# High-level stimulus template modulates neuronal response at the earlier processing stages

**DOI:** 10.1101/2020.04.14.041202

**Authors:** Vladimir Maksimenko, Alexander Kuc, Nikita Frolov, Semen Kurkin, Alexander Hramov

## Abstract

There is ample evidence that the brain matches sensory information with internal templates, but the details of this mechanism remain unknown. Here we consider the processing of repeatedly presented ambiguous stimuli with high ambiguity (HA) and low ambiguity (LA) and analyze how the processing depends on the ambiguity of the previous stimulus. On the behavioral level, we report a faster response to the HA stimulus after HA stimuli and a faster response to the LA stimulus after LA stimuli. The EEG analysis reveals that when HA stimulus follows LA stimuli, the neuronal activity in the sensory areas attenuates at the early processing stage but enhances during the latter stages. It evidences the hierarchical processing organization where low levels process the stimulus details, and high levels represent its interpretation. It also confirms that on low levels, HA and LA stimuli processing is similar due to their similar morphology. Therefore, the brain uses the LA stimulus template on the low levels to reduce the demands when processing the HA stimulus details. When LA stimulus follows HA stimuli, the attenuated response in the sensory regions accompanies high response in the frontal cortex. Namely, we observe high *θ* power in the medial frontal cortex and high *β* power in the right inferior frontal cortex. It shows activation of the top-down cognitive control functions detecting the mismatch between the LA stimulus and the HA stimulus template and transfer this template to the low processing levels.

**Significance statement:** The brain attenuates its response to repeatedly presented similar stimuli. When an ambiguous visual stimulus follows unambiguous stimuli with the same morphology, the neuronal response in sensory areas decreases at the early processing stage but enhances during the latter stages. It evidences hierarchical processing organization where low levels process the details, and high levels represent the interpretation. It also confirms that the brain uses templates on different levels to reduce the processing demands. When an unambiguous stimulus follows ambiguous stimuli, a low response in the sensory regions accompanies high response in the frontal cortex. It manifests activation of the top-down mechanisms to detect the mismatch between an unambiguous stimulus and an ambiguous template and transfer this template to low levels.

## Introduction

The brain uses sensory information to create a representation of the external environment. The growing body of literature suggests that information processing is organized hierarchically: low levels process detailed information, while high levels represent integrated information. There is a view that the brain uses the prior knowledge (predictions) along with the sensory evidence to create an accurate representation of the external environment (Rauss and Pourtois, 2013; Teufel and Fletcher, 2020).

The widespread tendency is to consider predictions as high-level processes acting in a top-down manner on mechanisms lower in the hierarchy. For instance, the bulk of literature suggests that top-down expectations lead to the formation of stimulus templates (Kok et al., 2014, 2017; Teufel et al., 2018). The brain matches these templates to external evidence (Heekeren et al., 2008): templates are transferred from high to low levels, whereas signals traveling in the opposite direction encode matching errors (Friston, 2009). Thus, the predictive signaling reflects top-down processes, and prediction-error signaling constitutes bottom-up processing. These processes are interdependent and always interact (Rauss and Pourtois, 2013). To date, there is ample evidence that the predictions are formed on the different processing levels and interact with each other via the top-down and bottom-up influences(Wacongne et al., 2011; Teufel and Fletcher, 2020). At the same time, the exact mechanism underlying the comparison between the predictions and the sensory evidence remain uncovered.

Here we consider the processing of repeatedly presented ambiguous stimuli, Necker cubes, with low ambiguity (LA) and high ambiguity (HA) degree. We instructed the subjects to report on each stimulus interpretation and recorded their EEG signals and the response time (RT). Repeated exposure to the same or similar sensory stimulus causes neuronal adaptation. This effect implies the reduced neural response to repeated compared with unrepeated stimuli (Henson and Rugg, 2003) and relates to the low-level (Vogels, 2016; Vinken et al., 2018) and the high-level (Gilbert and Li, 2013; Schwiedrzik and Freiwald, 2017) predictions. According to our previous works (Maksimenko et al., 2019, 2017), HA stimuli interpretation takes longer RT than the interpretation of LA ones. The LA and HA Necker cubes have almost the same morphology, and we assume their similar processing on low levels. We also suppose that the HA stimulus interpretation engages higher levels; therefore, on those levels, its processing requires a higher neuronal response. Finally, we suggest that the HA stimulus template appears on the hierarchically higher processing levels than the LA template. As a result, HA and LA templates may affect the ongoing stimulus processing in different ways.

To address this issue, we analyze the effect of the previous stimulus ambiguity on the processing of the ongoing visual stimulus. On the behavioral level, we demonstrate that the subjects respond faster to HA stimuli following HA stimuli. On the contrary, faster response to LA stimuli follows LA stimuli. The EEG analysis reveals that when HA stimulus follows LA stimuli, the neuronal activity in the sensory (occipito-parietal) areas attenuates at the early processing stage but enhances during the latter stages. It evidences the hierarchical processing organization where low levels process the stimulus details, and high levels represent its interpretation. It also confirms that on low levels, HA and LA stimulus processing is similar due to similar morphology. Therefore, the brain uses the LA stimulus template on the low levels to reduce the demands when processing the HA stimulus details. When LA stimulus follows HA stimuli, the attenuated neuronal response in the sensory regions accompanies high response in the frontal cortex. Namely, we observe the high *θ* power in the medial frontal cortex and the high *β*-band power in the right inferior frontal cortex. It manifests activation of the top-down cognitive control functions to detect the mismatch between the LA stimulus and the HA stimulus template and transfer this template to the low processing levels.

## Methods

### Participants

Twenty healthy subjects (11 males and 9 females) aged from 26 to 35 with normal or corrected-to-normal visual acuity participated in the experiments. All of them provided written informed consent in advance. All participants were familiar with the experimental task and did not participate in similar experiments in the last 6 months. The experimental studies were performed under the Declaration of Helsinki and approved by the local Research Ethics Committee of the Innopolis University.

## Experimental procedure

The Necker cube was used as an ambiguous visual stimulus (Kornmeier et al., 2017; Hramov et al., 2017). A subject without any perceptual abnormalities perceives the Necker cube as a 3D-object due to the specific position of the cube’s edges. Depending on the contrast of the inner edges, the Necker cube can be interpreted as either left- or right-oriented (See Fig. 1, a).

**Figure 1.**
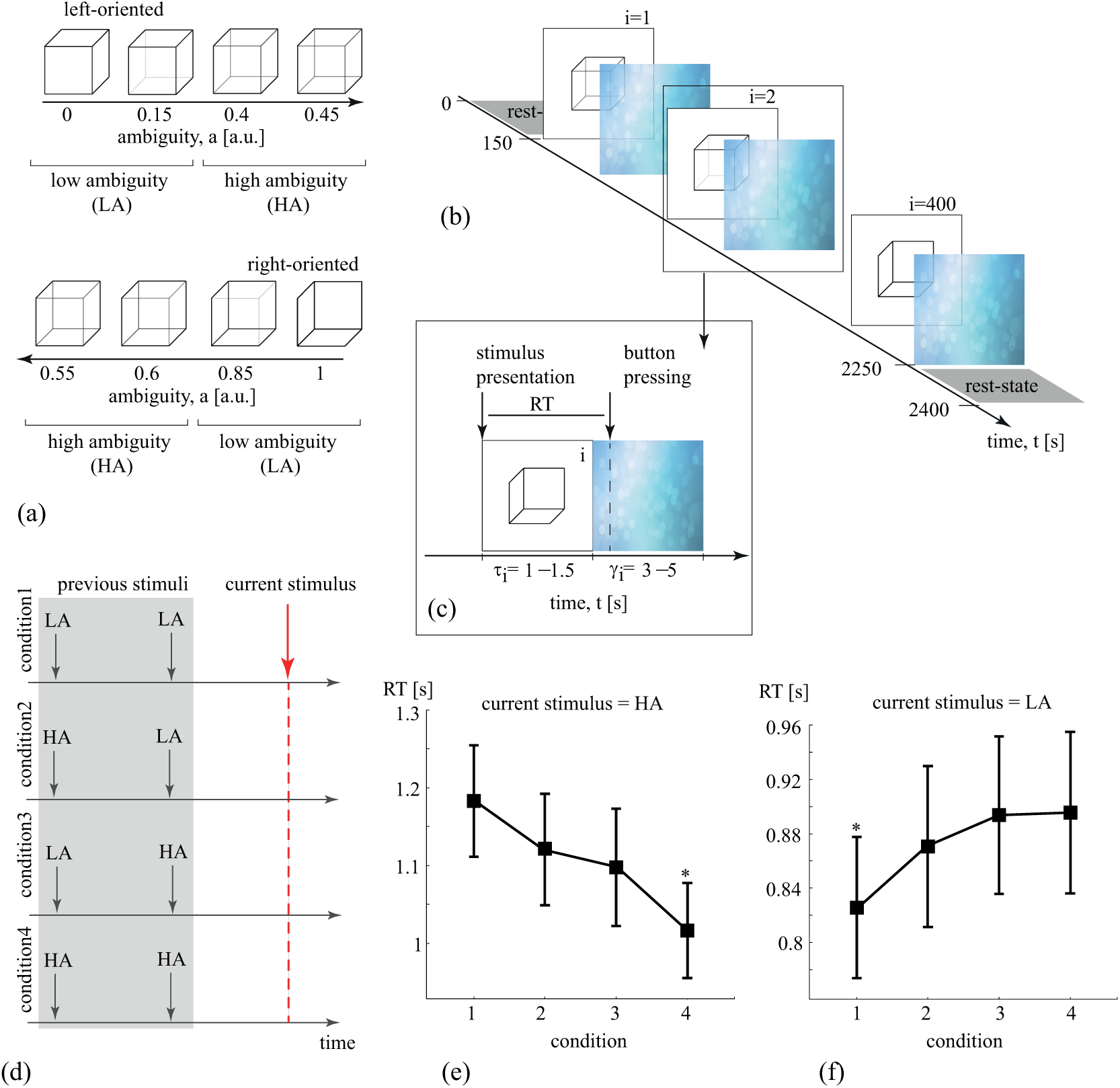
(a) The set of visual stimuli (Necker cubes) with a different ambiguity degree. (b) The structure of the experimental session. (c) The single image presentation: *τ_i_* is the stimulus presentation time, *γ_i_* is the abstract image presentation time, RT is the subject’s response time to the stimulus. (d) The four experimental conditions. (e) The RT of the HA stimulus processing in each condition (group mean SE). (f) The RT of the LA stimulus processing in each condition (group mean SE). *∗ p <* 0.05 via the Wilcoxon test, sample size *n* = 20 subjects.

The contrast of the three middle lines centered in the left middle corner was used as a control parameter *a* ∊ [0, 1]. The values *a* = 1 and *a* = 0 correspond, respectively, to 0 (black) and 255 (white) pixels’ luminance of the inner lines using the 8-bit gray-scale palette. Therefore, the control parameter can be defined as *a* = *g/*255, where *g* is the brightness of the inner lines. In our experiment, we use the Necker cube images with 8 different ambiguity levels (see Fig. 1, a). Half of them (*a* 0.0, 0.15, 0.4, 0.45) are considered as left-oriented and another half (*a* ∊ {0.55, 0.6, 0.85, 1}) as right-oriented. While for *a* ≈ 0 and *a* ≈ 1 (low ambiguous (LA) images) the cubes can easily be interpreted as left- and right-oriented, for *a* ≈ 0.5 the identification of the cube orientation is a more difficult task since we deal with highly ambiguous (HA) images.

Each Necker cube image was drawn by black and gray lines located at the center of the computer screen on a white background. The 14.2-cm Necker cubes were demonstrated on a 24” BenQ LCD monitor with a spatial resolution of 1920 × 1080 pixels and a 60-Hz refresh rate. The subjects were located at a 70–80 cm distance from the monitor with a visual angle of approximately 0.25 rad.

The whole experiment lasted around 40 min for each participant, including short recordings of the resting EEG state (≈ 150 ms) before and after the main part of the experiment. During experimental sessions, the cubes with predefined values of *a* (chosen from the set in Fig. 1, a) were randomly demonstrated 400 times, each cube with a particular ambiguity was presented about 50 times. The scheme of the experimental session is shown in Fig. 1, b. Each *i*-th stimulus was presented for time interval *τ_i_*, and the next (*i* + 1)-th image after time interval *γ_i_*.

The participants were instructed to press either the left or right key to indicate their first impression on the orientation of each successively presented cube. The consecutively presented images affect the perception of previously demonstrated cubes. For example, if the subject observed several left-oriented cubes in a row, then his/her perception was stabilized to the left-oriented cube, even if the next cube was right-oriented. Such memory is referred to as a stabilization effect (Leopold et al., 2002). In order to reduce this “memory” effect, the duration of the stimulus exhibition varied in the range of *τ* ∊ [1, 1.5] s. Moreover, a random variation of the control parameter *a* also prevented the perception stabilization. Lastly, to draw away the observer’s attention and make the perception of the next Necker cube independent of the previous one, different abstract pictures were exhibited for about *γ* ∊ [3, 5] s between subsequent demonstrations of the Necker cube images.

For each stimulus, we estimated a behavioral response by measuring the response time (RT), which corresponded to the time passed from the stimulus presentation to button pressing (Fig. 1, c).

### EEG recording and processing

The EEG signals were recorded using the monopolar registration method and the classical extended 10–10 electrode scheme. We recorded 31 signals with two reference electrodes A1 and A2 on the earlobes and a ground electrode N just above the forehead. The signals were acquired via the cup adhesive Ag/AgCl electrodes placed on the “Tien–20” paste (Weaver and Company, Colorado, USA). Immediately before the experiments started, we performed all necessary procedures to increase skin conductivity and reduce its resistance using the abrasive “NuPrep” gel (Weaver and Company, Colorado, USA). After the electrodes were installed, the impedance was monitored throughout the experiments. Usually, the impedance values varied within a 2–5 kΩ interval. The electroencephalograph “Encephalan-EEG-19/26” (Medicom MTD company, Taganrog, Russian Federation) with multiple EEG channels and a two-button input device (keypad) was used for amplification and analog-to-digital conversion of the EEG signals. This device possessed the registration certificate of the Federal Service for Supervision in Health Care No. FCP 2007/00124 of 07.11.2014 and the European Certificate CE 538571 of the British Standards Institute (BSI). The raw EEG signals were filtered by a band-pass FIR filter with cut-off points at 1 Hz (HP) and 100 Hz (LP) and by a 50-Hz notch filter by embedded a hardware-software data acquisition complex. Eyes blinking and heartbeat artifact removal was performed by Independent Component Analysis (ICA) using EEGLAB software (Delorme and Makeig, 2004). After the EEG preprocessing procedure, we excluded some trials due to the high-amplitude artifacts and considered 320 trials out of the initial 400.

The recorded EEG signals were segmented into 4-s trials, where each trial was associated with a single presentation of the Necker cube, including 2-s interval before and 2-s interval after the moment of the Necker cube demonstration. We calculated the wavelet power for each trial in the 4−40 Hz frequency band using the Morlet wavelet. The number of cycles (*n*) was defined as *n* = *f*, where *f* is the signal frequency. The wavelet analysis was performed in Matlab using the Fieldtrip toolbox. The 0.5 s intervals on each side of the trial were reserved for the wavelet power calculation. As a result, we considered the wavelet power on the 3 s interval including the prestimulus baseline (from −1.5 to 0 s) and the stimulus-related activity (from 0 s to 1.5 s). For the stimulus-related wavelet power we performed the baseline correction as [stimulus-related activity–prestimulus baseline]/prestimulus baseline.

### Behavioral estimates and the condition selection

First, all Necker cubes were divided into 4 groups according to their ambiguity and orientation: the left-oriented cubes with the low ambiguity (*a* ∊ 0, 0.15); the left-oriented cubes with the high-ambiguity (*a* ∊ 0.4, 0.45); the right-oriented cubes with the low-ambiguity (*a* ∊ 0.85, 1); the right-oriented cubes with the high-ambiguity (*a* ∊ 0.55, 0.6). For each group, we divided the RT distribution into two sets of trials: low RT trials lying between the 2.5 percentile and the median; high RT trials lying between the median and 97.5 percentile. As the result, the following experimental conditions were considered: *ambiguity* (LA vs HA), *orientation* (left vs right) and *RT* (low vs high). The number of trials was kept constant for all conditions and all subjects. As a result, each condition included 35 trials.

Second, we grouped LA and HA cubes according to the ambiguity of the previous cube. We considered not only the previous (1-back) cube ambiguity, but the ambiguity of the cube presented two turns back (2-back). As a result, for LA and HA cubes, we considered four conditions (See Fig. 1, d): *condition*1 – both previous cubes have low ambiguity (LA-LA); *condition*2 – the 1-back cube has the low ambiguity, and the 2-back cube has the high ambiguity (HA-LA); *condition*3 – the 1-back cube has the high ambiguity, and the 2-back cube has the low ambiguity (LA-HA); *condition*4 – both previous cubes have high ambiguity (HA-HA). To keep the number of EEG trials constant across the conditions and subjects, we considered 16 trials for each condition.

### Statistical analysis

The statistics on the group level was performed for the median RT, median presentation time and the ratio between the left- and right oriented cubes and the ratio between the LA and HA cubes. The main effects were evaluated via a repeated measures ANOVA. The Greenhouse-Geisser correction was used depending on the results of the spherisity tests. The post hoc analysis was performed either via the paired samples t-test or via the Wilcoxon signed rank test depending on the samples normality. Normality was tested via the the Shapiro-Wilk test. The statistical analysis was carried out in SPSS. The used tests as well as their parameters are mentioned in the results section.

The wavelet power topograms were compared for the different experimental conditions in the time, spatial and frequency domain via the cluster-based permutation test. The critical *α*-level for the pairwise comparison was set to 0.05, and the critical *α* level for the cluster-level statistics was set to 0.025 corresponding to a false alarm rate of 0.05 in a two-sided test. Finally, the minimal number of the elements in the cluster was set to 2, and the number of permutations was equal to 2000. The topograms were compared using the Fieldtrip toolbox in Matlab.

### Source localization

We used the exact low-resolution brain electromagnetic tomography (eLORETA) method to solve the inverse problem and reconstruct source activity from the EEG at each of the predefined points over brain volume (Pascual-Marqui, 2007; Pascual-Marqui et al., 2006; Pascual-Marqui, 2009). eLORETA is a linear, regularized, weighted minimum norm inverse solution with theoretically exact zero error localization even in the presence of structured biological or measurement noise (Pascual-Marqui, 2007; Grech et al., 2008).

“Colin27” brain MRI averaged template (Holmes et al., 1998) was used to develop a boundary element method (BEM) head model with three layers (brain, skull, and scalp) (Fuchs et al., 2002; Baillet et al., 2001) and source space consisting of 24570 voxels in the brain. The EEG electrodes’ positions were fitted to the template head shape. The use of template models has previously been demonstrated to perform well compared to individual models derived from MRI (Fuchs et al., 2002; Lopes et al., 2020).

All operations were performed using the Fieldtrip toolbox (Oostenveld et al., 2011). The inverse solution yielded estimates of power of sources at each voxel averaged over the chosen time window.

Sources reconstruction was carried out separately for each subject and each condition. At first, every trial was demeaned and filtered by the fourth-order Butterworth bandpass filter with the passband of [*f_L_, f_H_*]. Then, we performed averaging across the trial belonging to each condition.

Since *t* = 2 s is the moment of the stimulus presentation, [1.9 – *dt*; 1.9] s was chosen as the pre-stimulus segment, and [2; 2 + *dt*] s — as the post-stimulus one. The sources were reconstructed separately for the pre-stimulus and post-stimulus segments. For these segments we obtained the averaged distributions of sources power over the brain volume (*P_pre_* and *P_post_*). We contrasted the post-stimulus power with the baseline in the following way: *P_con_* = (*P_post_ P_pre_*) */P_pre_*.

The filter band *f_L_*, *f_H_* and *dt* were chosen depending on the frequencies of interest: *f_L_* = 5 Hz, *f_H_* = 7 Hz, *dt* = 0.2 s – for the *θ*-band, and *f_L_* = 18 Hz, *f_H_* = 20 Hz, *dt* = 0.1 s – for the *β*-band.

Finally, for the *θ* - and the *β*-bands, we obtained the source power distributions in two conditions. Each condition included 20 distributions (every distribution corresponded to one subject). To determine the significant difference between the conditions, we conducted a statistical cluster-based permutation test (Maris, 2012; Maris and Oostenveld, 2007). To match the discovered significant clusters with the anatomical regions of the brain, we used the Automated Anatomical Labeling (AAL) brain atlas (Tzourio-Mazoyer et al., 2002).

## Results

### Effect of the previous stimulus orientation and ambiguity

First, we consider whether the properties of the previous cube (orientation and ambiguity) affect RT for the current cube. We introduce three within-subject parameters which characterize the current cube: *ambiguity* (LA vs HA), *orientation* (left vs right) and *RT* (low vs high) (see Methods). We perform a repeated-measures ANOVA to test how the ratio between the number of the left and the right-oriented previous cubes 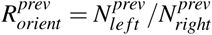 and the ratio between the number of LA and HA previous cubes 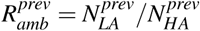 differs between the different current cube’s *RT* depending on its *orientation* and *ambiguity*.

As a result, ANOVA with the Greenhouse-Geisser correction reveals that 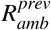 differs insignificantly between low and high RT (*F*_1,19_ = 0.004*, p* = 0.952). At the same time, the significant interaction effect (*RT* × *ambiguity*) is observed (*F*_1,19_ = 11.89, *p* = 0.003) while the interaction effect (*RT* × *orientation*) (*F*_1,19_ = 0.011*, p* = 0.918) and (*RT* × *ambiguity×orientation*) (*F*_1,19_ = 0.008*, p* = 0.93) are insignificant. These results mean that the ratio between the number of LA and HA previous cubes affects the RT in different ways, depending on the current cube ambiguity. We perform the post-hoc analysis to compare 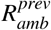 separately for HA and LA current cubes. The samples obey the Shapiro-Wilk normality test and have been compared via a paired samples *t*-test. As the result, the significant increase of 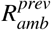 has been observed for LA cubes when compared high RT with low RT (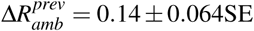, *t* = 2.242, *d f* = 19, *p* = 037). On the contrary, for HA cubes the 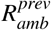 is lower for high RT when compared with low RT (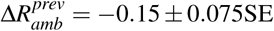, *t* = 2.032, *d f* = 19, *p* = 0.056). The results evidence that the RT for the current cube is lower if the previous cube has the same ambiguity. Note that the observed effect is significant for the LA cubes, while for HA cubes, the reported *p*-value is close to the critical.

For the 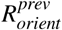 ANOVA with the Greenhouse-Geisser correction reveals significant interaction effect (*RT × ambiguity× orientation*) (*F*_1,19_ = 5.430*, p* = 0.031), while the interactions (*RT × ambiguity*) (*F*_1,19_ = 0.788*, p* = 0.386) and (*RT × orientation*) (*F*_1,19_ = 0.384*, p* = 0.543) were insignificant. Finally, no significant difference of 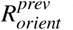 has been observed for high and low RT in general (*F*_1,19_ = 0.767*, p* = 0.392). Thus we conclude that the previous cube orientation affects the RT of the current cube in different ways depending on the ambiguity and the orientation of the current cube. Therefore, we perform the post hoc analysis to compare 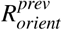 for the high and low RT for the different ambiguity and the different orientations of the current cube. The post hoc analysis with the paired samples t-test reveals the significant increase of the 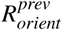 for the left-oriented LA cubes when compared high RT to the low RT (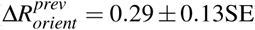, *t* = *−*2.254, *d f* = 19, *p* = 0.036). In the rest cases, 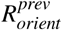 differs insignificantly between high and low RT. These results demonstrate that the orientation of the previous cubes affects the RT only in the case of the left-oriented LA cubes. The RT for the left-oriented LA cubes is lower if the previous cube is right-oriented.

### Previous stimulus ambiguity affects the response time

In further analysis, we focus on the effect of the previous cube ambiguity separately for the LA and HA current cubes. To analyze this effect more in detail, we consider not only the previous (1-back) cube ambiguity, but the ambiguity of the cube presented two turns back (2-back) and compare RT in the four conditions (see Methods and Fig. 1, d for the details). The repeated measures ANOVA reveals the significant difference of RT between the conditions for both HA (*F*_3,57_ = 10.787*, p <* 0.001) and LA (*F*_3,57_ = 6.067*, p* = 0.001) current cubes. Fig. 1 demonstrates how the RT (group mean ±SE) differs in these conditions for HA (e) and LA (f) cubes.

For the HA cubes, we observe the minimal RT in the *condition*4 where the two previous cubes have the high ambiguity and the maximal RT – in the *condition*1 where the two previous cubes have the low ambiguity. The post hoc analysis with the Wilcoxon test reveals that the RT in the *condition*4 is significantly lower when compared to the *condition*1 (*Z* = 3.547*, p <* 0.001), *condition*2 (*Z* = 3.323*, p* = 0.001) and *condition*3 (*Z* = 2.696*, p* = 0.007).

For the LA cubes, we observe the minimal RT in the *condition*1 where the two previous cubes have the low ambiguity and the maximal RT- in the *condition*4 where the two previous cubes have the high ambiguity. The Wilcoxon test reveals the RT in the *condition*1 is significantly lower when compared to the *condition*2 (*Z* = 2.67*, p* = 0.008), *condition*3 (*Z* = 2.13*, p* = 0.009) and *condition*4 (*Z* = 3.061*, p* = 0.002).

Finally, ANOVA with the Greenhouse-Geisser correction reveals that the median presentation time does not significantly change across the conditions for both LA (*F*_1.315,24.991_ = 2.883*, p* = 0.093) and HA (*F*_1.387,26.354_ = 1.646*, p* = 0.214) current cubes. The ratio between the number of the left and right orientation of the current cube also differs insignificantly across the conditions for both LA (*F*_1.262,29.020_ = 3.604*, p* = 0.059) and HA (*F*_1.981,45.55_ = 1.159*, p* = 0.323) cubes. These results evidence that neither the time spent in the experiment nor the orientation but the previous cube ambiguity affects RT in the considered conditions.

### Neural activity during prestimulus state

According to the results described above, the ambiguity of the previous cube affects RT for the current cube. Thus RT is lower if the previous cube ambiguity coincides with the current cube ambiguity. This effect is stronger if two previously presented cubes have similar ambiguity (both LA or both HA). We suppose that the processing of the previous cubes affects the properties of the prestimulus state. The prestimulus state following the LA cubes processing (condition1) is more beneficial for the processing of the ongoing LA cube. In contrast, the prestimulus state following the HA cubes processing is more beneficial for the processing of the ongoing HA cube. According to this, we compare the prestimulus states corresponding to the condition1 and condition4 (Fig. 2, a).

**Figure 2.**
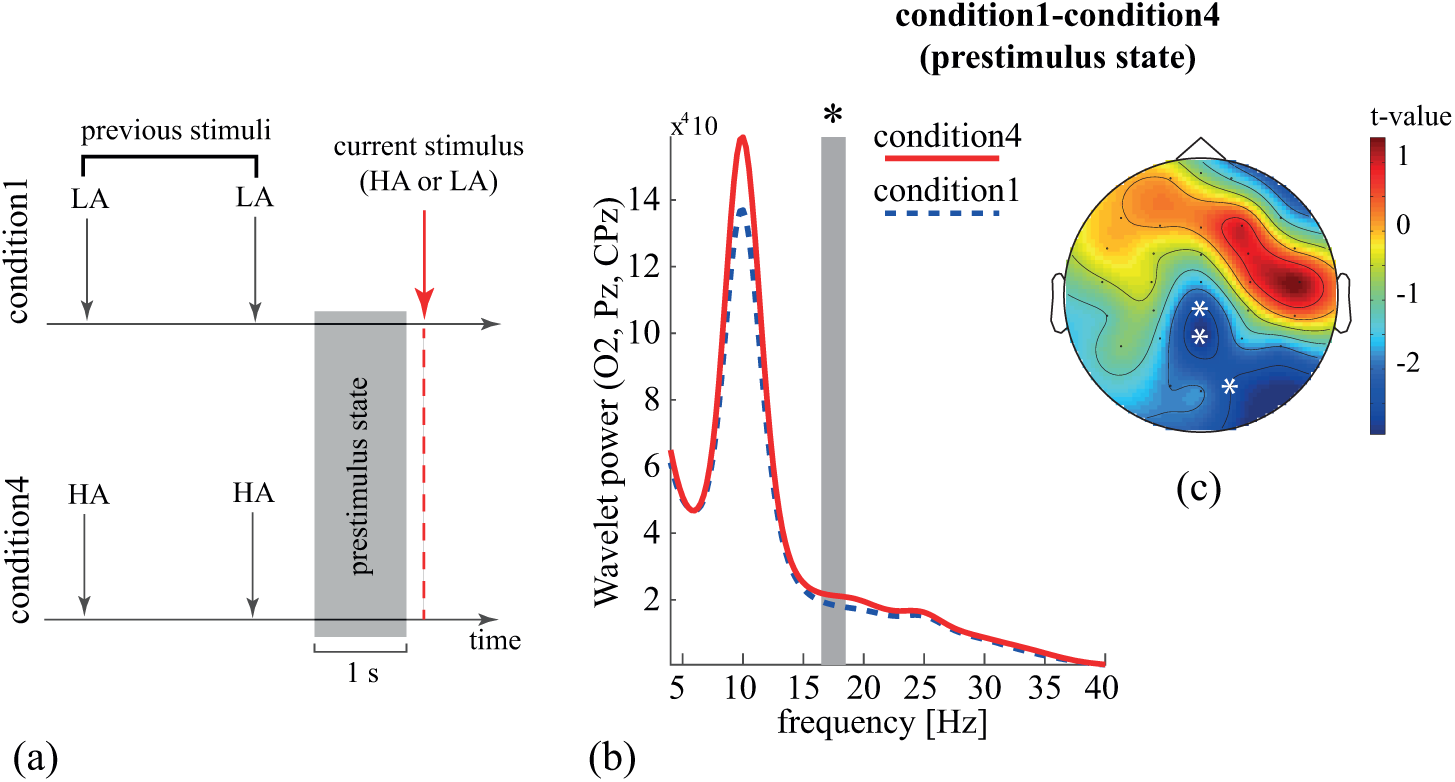
The RT for HA (a) and LA (b) Necker cube in four different experimental conditions (group mean, SD and individual values). Each experimental condition represents the different ambiguity of two previously presented Necker cubes. *∗ p <* 0.01 via the Wilcoxon test.

Here we do not consider the ambiguity of the current cube as a within-subject factor. Therefore, the power spectra have been averaged across the HA and LA current cubes. As a result, the only one negative cluster has been observed in the *β*-frequency range (Fig. 2, b) indicating the higher wavelet power in condition 4. The comparison of the topograms reveals that this cluster is localized in the occipito-parietal area (Fig. 2, c). Note that the median presentation time of the current cube does not significantly differ for these conditions (*Z* = 1.307*, p* = 0.191, Wilcoxon test) as well as the ratio between the left- and right-oriented cubes (*Z* = 0.105*, p* = 0.917, Wilcoxon test). Thus we conclude that the observed cluster is caused by only the previous cubes ambiguity but not the experiment duration or the cube orientation.

### Neural activity during HA stimuli processing

As described above, after the processing of two HA cubes, the brain state is characterized by the increased *β*-band power over the occipito-parietal EEG sensors. In this case, the processing of the ongoing HA cube takes the lowest RT (see Fig. 1, e). To analyze the cortical activity underlying the decreased RT in this condition, we compare the wavelet power in the condition 1 and condition 4 in the time (0.35 s after the stimulus presentation) and frequency (4-40 Hz) domain. As a result, two positive clusters and no negative clusters have been found. The clusters’ location in the time-frequency area are shown in Fig. 3, a. One can see that the *β*-cluster (I) appears 0.1 s after the stimulus presentation. It includes the occipital and parietal EEG sensors in the right hemisphere (O2, P4, P8, TP8) (see Fig. 3, b). The *θ*-cluster (II) is observed for 0.1-0.25 s after the cube presentation. According to Fig. 3, c, it includes the parietal (P3, Pz, P4) and occipital (O1) sensors.

**Figure 3.**
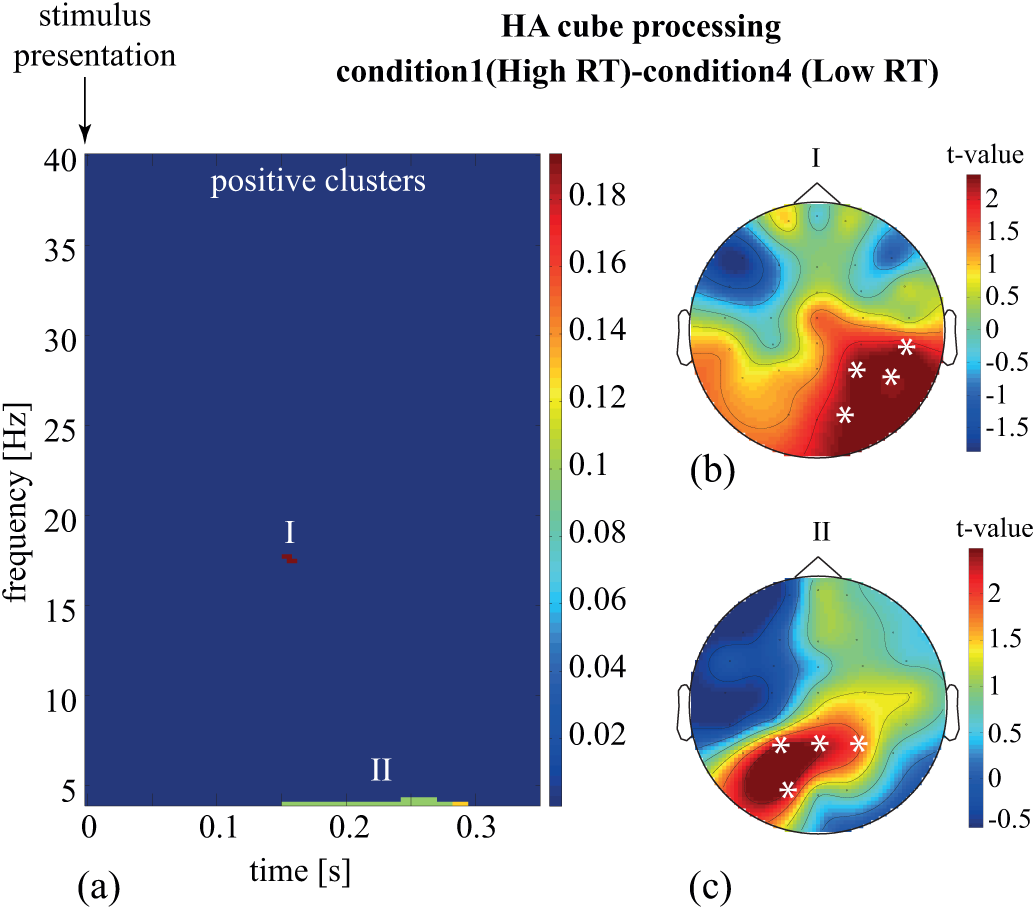
The EEG channel clusters representing the significant positive change of the spectral energy between condition1 (LA-LA) and condition4 (HA-HA) during the HA cube processing: clusters location in the time-frequency domain (a) and spatial configuration (b-c).

### Neural activity during LA stimuli processing

When comparing the cortical activity during the LA processing in condition 1 and condition 4, we have found four negative and three positive clusters. The negative clusters are observed during the first 0.15 s after the stimulus presentation. On can see the *β* cluster (I) and three *θ* clusters (II-IV) (Fig. 4, a). The *β* cluster (I) locates in the right temporal and frontal areas (Fig. 4,b). The *θ* clusters (II-IV) belong to a single form of the *θ* activity in the left parietal (P3, CPz), temporal (T5, FT7) and the midline frontal (FCz, Fz) electrodes (Fig. 4, c-e). The positive clusters appear in *β* frequency band 0.2 s after the stimulus presentation (see Fig. 6, a). The *β*-cluster (I) is observed for ~ 17 Hz for 0.2 s in right parietal (P4), frontal (FC4) and temporal sensors (Fig. 6, b). Another *β* cluster (II) appears after 0.25 s in the 20−25 Hz band in the right occipital (Oz, O2), parietal (P4) and temporal (P8, TP8) electrodes (Fig. 6, c). Finally, the *β*-cluster (III) is observed after 0.3 s in the 25−30 Hz frequency band. This cluster includes occipital (Oz, O2), parietal (P4, P8), central (Cz) sensors mostly in the right hemisphere (Fig. 6, d).

**Figure 4.**
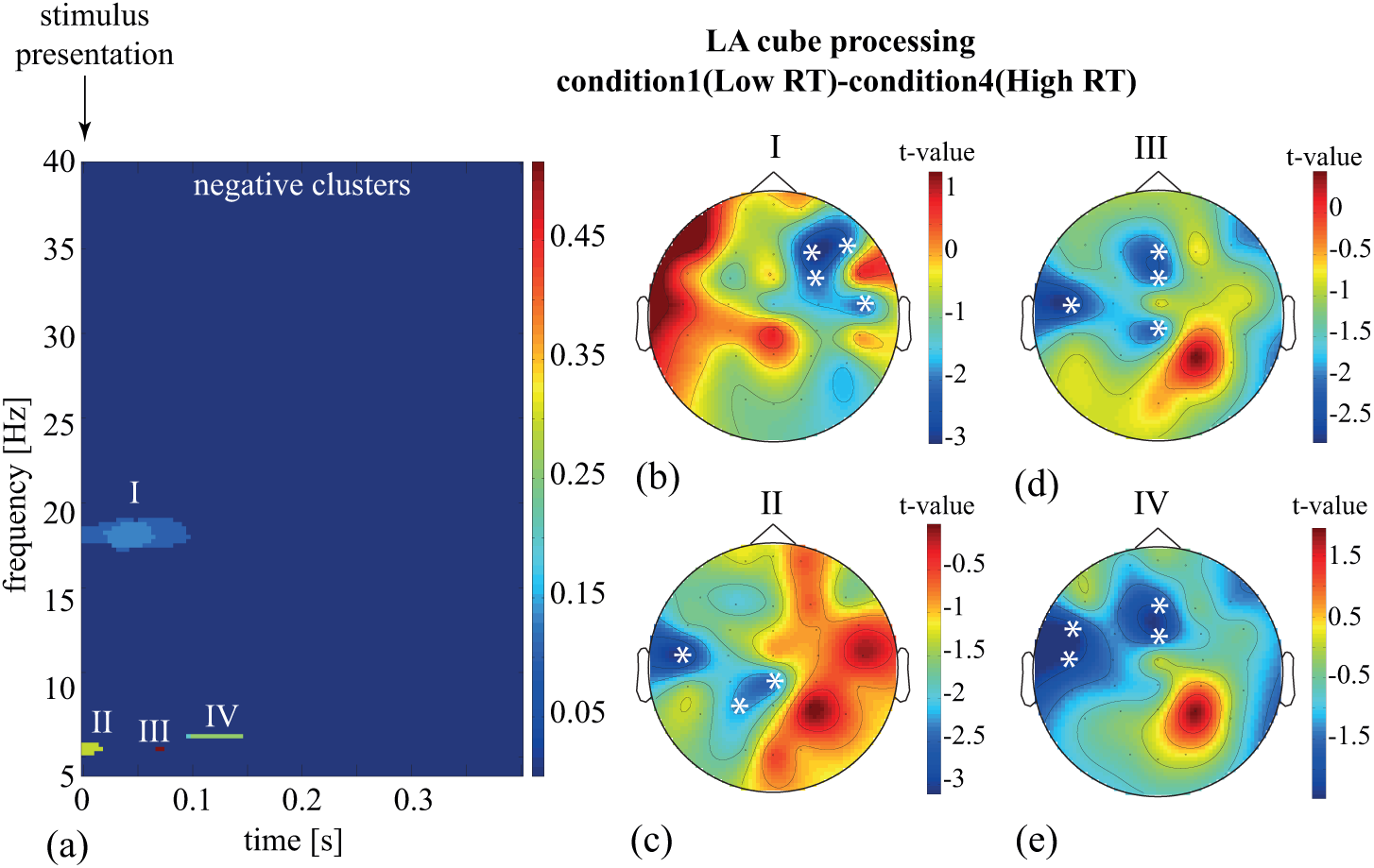
The EEG channel clusters representing the significant negative change of the spectral energy between condition1 (LA-LA) and condition4 (HA-HA) during the LA cube processing: clusters location in the time-frequency domain (a) and spatial configuration (b-e).

**Figure 5.**
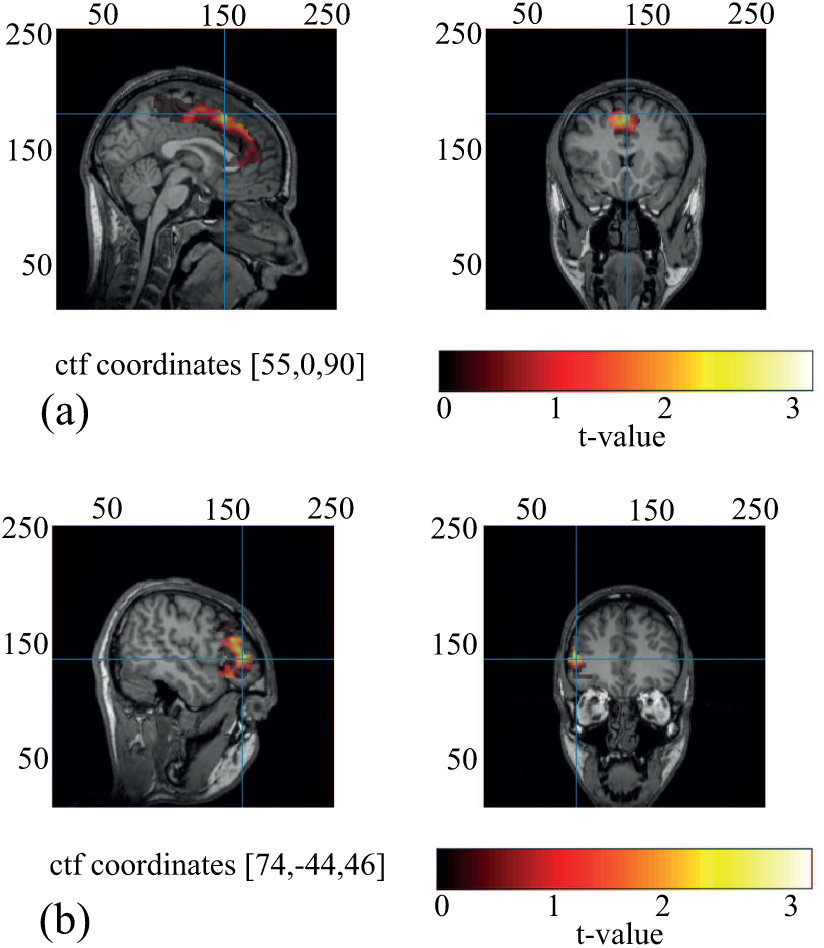
The frontal *θ* (a) and *β* (b) clusters in the source space. The color scale represents *t*-value as the result of the comparison between condition1 and condition4. The crossing lines reflect the location of the cluster (*p_pairwise_ <* 0.01, *p_cluster_ <* 0.025). *t*-values are shown for the cingulum (a) and right inferior frontal gyrus (b)

**Figure 6.**
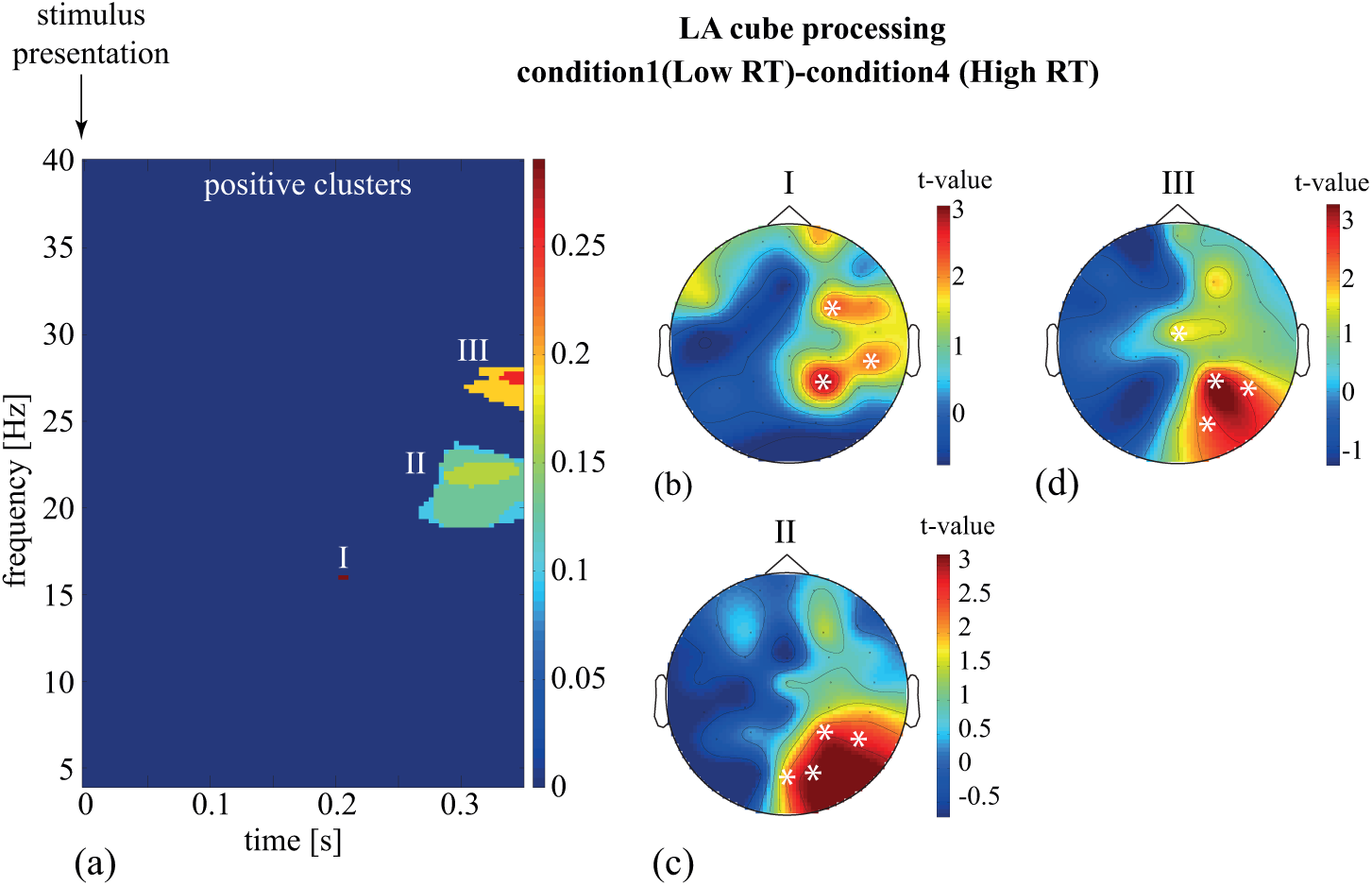
The EEG channel clusters representing the significant positive change of the spectral energy between condition1 (LA-LA) and condition4 (HA-HA) during the LA cube processing: clusters location in the time-frequency domain (a) and spatial configuration (b-d).

The *β* and *θ* clusters, which appear during the LA cube processing in the condition4, have been localized in the source space. We have shown that the source of the *θ* activity belongs to the medial cingulate cortex (Fig. 5, a), whereas the source of the *β* activity appears in the right inferior frontal cortex (Fig. 5, b).

## Discussion

When perceiving the subsequently presented ambiguous stimuli, Necker cubes, the previous stimulus ambiguity affects the current stimulus processing. We show that the subject responds faster to the stimulus if its ambiguity matches the ambiguity of the previously presented stimuli.

The reasonable explanation is that the neuronal populations involved in the processing of the previous stimuli remain active during the the following prestimulus period. Having compared the prestimulus period following HA and LA stimuli processing, we have found the increased occipito-parietal *β*-band power in the former case. The activation of neuronal populations before the stimulus processing can be related to the prestimulus template (Kok et al., 2014, 2017).

According to (Summerfield and De Lange, 2014), the stimulus template appears as the result of the top-down expectations. If so, to respond faster, the subjects expect HA stimulus after two HA stimuli. However, we present Necker cubes randomly, and the probability of seeing HA cube after two HA cubes is small. The subjects are more likely to expect LA stimulus after two HA ones. Finally, they have to report the particular orientation of the Necker cube, but not its ambiguity. Therefore, the expectation and the resulting template should include the stimulus orientation. Given the above, the subjects should respond faster to the stimulus with the same orientation as the previous one. Our results, in turn, demonstrate that a similar orientation of the current and the previous stimuli do not cause a faster response.

According to the review (Rauss and Pourtois, 2013), the stimulus templates appear on the different levels of the sensory processing hierarchy. We suppose that the stimulus orientation characterizes the high-level template, whereas the template on the lower levels reflects the stimulus morphology. The neuronal response during the HA stimulus processing increases due to the two possible reasons. First, HA stimulus has an uncertain orientation, thus to determine it, the lager neuronal populations are engaged on the high levels of the processing hierarchy. Second, HA stimulus morphology is more complicated. Along with the similar features (outer edges), the HA Necker cube has a higher contrast of the inner edges. Thus, to process the HA stimulus, the more substantial amount of the sensory evidence has to be acquired, resulting in the higher neuronal activation on the low levels of the processing hierarchy. Since the orientation of the previous stimulus does not affect the current stimulus processing, we do not consider the high-level template. On the contrary, we focus on the low-level templates.

We have compared the stimulus-related activity during the HA stimulus processing between two cases: the previous LA and the previous HA stimuli. We have observed the higher *β* - and *θ*-band power across the occipital and parietal sensors in the former case for *t <* 0.15 s. The increased parietal *θ*-band power evidences the increased processing demands resulting in mobilization of the cognitive resources, i.e., the top-down attentional control and the working memory (Tseng et al., 2018; Berger et al., 2019). The high occipital and parietal *β*-band power characterizes the tasks that involve endogenous top-down processes, including the ambiguous stimuli processing (Engel and Fries, 2010). In particular, in Ref. (Okazaki et al., 2008), the bistable visual stimuli (face-saxophone) were presented to the subjects after the pictures that were clear representations of the face or the saxophone. Similarly to our results, the increased *β* band power was observed in the occipital and the parietal regions during the ambiguous stimulus processing following the unambiguous one. The authors reported that the subjects mostly interpreted an ambiguous picture as the face. The *β*-band power was higher if the ambiguous figure followed the saxophone rather than the face image. Since the unambiguous stimulus was a part of the ambiguous one, their morphology was assumed to be similar. Thus authors suggested that the enhanced neuronal response was associated with the high-level processes related to the stimulus interpretation as a whole rather than with the low-level processing of the particular stimulus features. As mentioned above, we do not consider the effect of the stimulus orientation. Therefore the higher processing stages responsible for the stimulus interpretation as a whole were excluded from the analysis. We concluded that higher *θ* and *β*-band power might be associated with the processing of the HA stimulus features, which are not similar to LA stimuli (e.g., the inner edges). The fact that we observe the higher response for *t >* 0.15 s after the stimulus presentation support this suggestion. Indeed, *t <* 0.15 s can be associated with the low levels where the outer contours of the Necker cube are processed. These stimulus features are similar for HA and LA cubes, and therefore, we assume the similar neuronal activity on these processing levels for LA and HA stimulus.

We have compared the stimulus-related activity during the LA stimulus processing between two cases: the previous LA and the previous HA stimuli. Similarly, the neuronal response in the occipital and parietal regions remains unchanged for *t <* 0.15 s. For *t >* 0.15 s, we observe the increased parietal *β*-band power for the previous LA stimulus. It supports our proposal that LA and HA stimuli are processed similarly at the earlier stages, whereas higher stages of HA stimulus processing requires additional resources. An unexpected finding is the higher *β* - and *θ*-band power observed for *t <* 0.15 s for the previous HA stimulus. The frontal-midline *θ*-band power increases when LA stimulus follows the HA one. We have found the source of the *θ*-power in the anterior cingulate cortex. According to the Refs. (Enriquez-Geppert et al., 2014; Cavanagh and Frank, 2014), the frontal-midline *θ* is a candidate biophysical mechanism for cognitive control. Cognitive control enables monitoring and detecting cognitive interference in the stream of sensory information processing (Botvinick et al., 2004). The ERP studies report that a fronto-medial scalp ERP amplitude grows in interference situations requiring more cognitive control. A recent review (Folstein and Van Petten, 2008) relates the fronto-medial ERP amplitude with two subcomponents of the cognitive control – detection of novelty or mismatch from a perceptual template and the response inhibition. Finally, in the Ref. (Nigbur et al., 2011), the frontal-midline *θ* power is shown to increase during the tasks accomplishing in the framework of the classical interference paradigms. The *β*-band power in the right frontal cortex is usually associated with the top-down inhibitory control (Wagner et al., 2018). This cognitive function is mobilized in the tasks that require a rapid stopping of action. For instance, in the stop-signal task, the participants are instructed to initiate a Go response on each trial, which they sometimes have to try to stop when a stop signal is subsequently presented. The recent study (Wagner et al., 2018) reports that a brain signature for action stopping is related with a right frontal *β* power increases following the stop signal but before the time of stopping. The authors further show that the same frontal right-lateralized beta signature of stopping is active after the unexpected events.

The absence of the differences in the occipital and parietal cortex at the earlier stages (*t <* 0.15 s) could indicate that the brain has formed appropriative templates on the low processing levels regardless of the previous stimulus ambiguity. It can be the result of the similar morphology of the outer edges of the HA and LA cubes. At the same time, activation of the cognitive control may reflect the significant initial mismatch between the HA cube template and the sensory input at the earlier processing stages. To explain this phenomenon, we assume that the HA stimulus is finally processed on the higher levels when compared to LA stimulus. Therefore, the HA template appears on higher levels than the LA template. The brain matches the incoming sensory information with the high-level stimulus template to minimize the processing costs. The sensory evidence on the low levels is insufficient to fit this template, and the brain tries to construct the low-level templates form the high-level one. This mechanism goes through the interaction between the top-down processes, related to the template adaptation and the bottom-up processes associated with the matching errors. Thus, when LA stimulus follows HA one, the more top-down control is required to translate the high-level HA template to the low processing levels. This hypothesis coincides with the view proposed by Rauss and Pourtois in Ref. (Rauss and Pourtois, 2013). The authors assumed that the higher levels of the processing hierarchy extract the high-order stimulus features and use them to create a set of predictions that can be rapidly adapted or exchanged. They also suppose that these predictions are transferred to the low processing levels via direct connections. According to these assumptions, authors relate these direct connections with the top-down processes, which enables information transfer between the high and low levels and skip the intermediate levels. Our results suggest that these top-down processes include conflict control and the inhibitory control subserved by the neuronal activity in the medio-frontal cortex, anterior cingulate cortex, and the right inferior frontal cortex.

## Conflict of interest statement

The authors declare that they have no competing interests

### Acknowledgments

This study is supported by the President Program (NSh-2594.2020.2) in the part of the experimental work. VM and AK are supported by the President Program (grant MK-1760.2020.2) in the part of the behavioral and EEG data analysis. SK is supported by the President Program (MD-1921.2020.9) in the part of the source reconstruction.

## Author contributions statement

VM and AH designed and supervised the study, AK and NF performed the data analysis on the sensor level, VM analyzed behavioral data, SK performed the data analysis on the source level. VM and NF wrote the Manuscript.

